# Long-term research needed to avoid spurious and misleading trends in sustainability attributes of no-till

**DOI:** 10.1101/788240

**Authors:** Sarah Cusser, Christie Bahlai, Scott M. Swinton, G. Philip Robertson, Nick M. Haddad

**Affiliations:** W.K. Kellogg Biological Station, Department of Integrative Biology, Michigan State University, 3700 East Gull Lake Dr., Hickory Corners, MI 49060, U.S; Department of Biological Science, Kent State University, 249 Cunningham Hall, Kent, OH 44240, U.S; Department of Agricultural, Food, and Resource Economics, Michigan State University, East Lansing, MI 48824, U.S; Great Lakes Bioenergy Research Center, Michigan State University, East Lansing, MI 48824, U.S; Department of Plant, Soil, and Microbial Sciences, Michigan State University, East Lansing, MI 48824, U.S

**Keywords:** yield, soil moisture, N_2_O fluxes, LTER, power analysis, moving window, profitability of no-till adoption

## Abstract

Agricultural management recommendations based on short-term studies can produce findings inconsistent with long-term reality. Here, we test the long-term relative profitability and environmental sustainability of continuous no-till agriculture practices on crop yield, soil moisture, and N_2_O fluxes. Using a moving window approach, we investigate the development and stability of several attributes of continuous no-till as compared to conventional till agriculture over a 29-year period at a site in the upper Midwest, U.S. We find that over a decade is needed to detect the consistent benefits of no-till on important attributes at this site. Both crop yield and soil moisture required periods 15 years or longer to generate patterns consistent with 29-year trends. Only marginally significant trends for N_2_O fluxes appeared in this period. Importantly, significant but misleading short-term trends appeared in more than 20% of the periods examined. Relative profitability analysis suggests that 10 years after initial implementation, 86% of periods recuperated the initial expense of no-till implementation, with the probability of higher relative profit increasing with longevity. Results underscore the essential importance of decade and longer studies for revealing the long-term dynamics and emergent outcomes of agricultural practices for different sustainability attributes and are consistent with recommendations to support the long-term adoption of no-till management.

**GRAPHICAL ABSTRACT:** 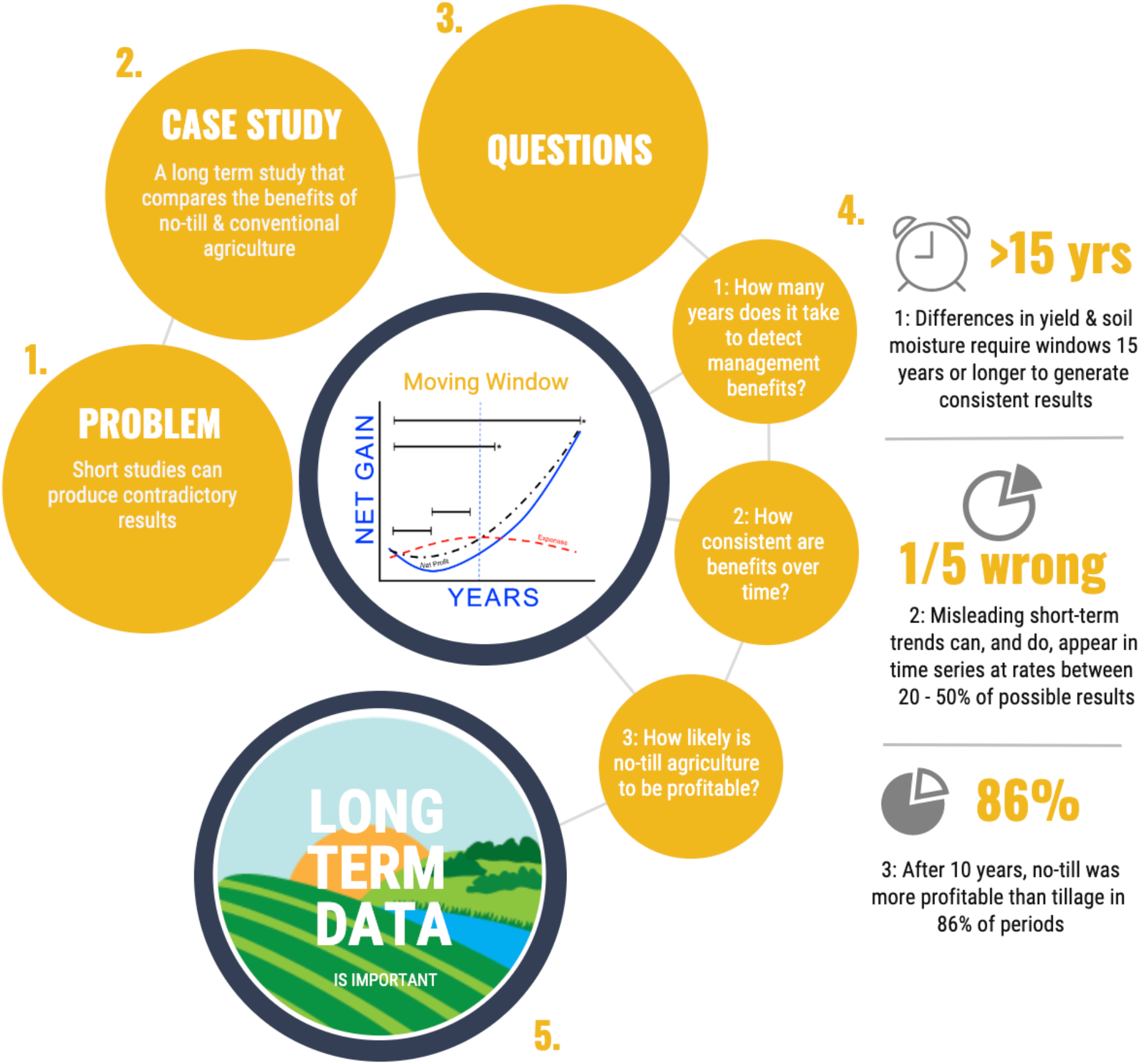

**HIGHLIGHTS:**

1. We test long-term effects of no-till on yield, soil moisture, and N_2_O fluxes
2. We examine 29 years of data with a moving window and relative profitability method
3. It takes at least a decade to detect consistent benefits of no-till
4. Shorter studies can produce significant but misleading findings
5. Long studies are essential to reveal the dynamics of agricultural management

## 1 INTRODUCTION

Agricultural decisions must promote sustainable benefits in the long-term. Considerable investment is made to agricultural research and development to inform these important land management decisions (USDA, 2007). The U.S. alone invests between $4.6 and $8.8 billion each year in public and private agricultural research (USDA, 2007) to promote innovative advances for many aspects of cropping system management (NRC, 2003; Robertson et al. 2008; Spiegal et al., 2018), including crop fertility, pest protection, seed genetics, water management, and soil health, among others. However, the outcomes of many management changes can be slow to develop and detect, especially those that depend on slow-to-change attributes of a system such as soil structure and organic matter. This adds to uncertainty regarding the long-term sustainability of management decisions (Robertson et al., 2008), raising the potential for misinterpretation of short-term studies.

Tillage is a well-researched management strategy that is a case in point. Tillage mechanically incorporates crop residues and/or amendments into soils, controls weeds, promotes soil warming and drying, and thereby prepares soil for planting, creating optimal conditions for crop seed germination and emergence (Reicosky et al., 1995). However, the environmental costs of regular tillage are great, including decreased soil carbon, increased potential for soil erosion, and poor soil structure (Lal et al., 2007b). Following the 1930’s dustbowl, U.S. agronomists and soil scientists became increasingly concerned about the long-term environmental sustainability of tillage-based agriculture (Derpsch et al., 2010). With the advent of modern herbicides in the 1960’s, along with glyphosate resistant crops in the 1990’s, no-till, whereby crop residues are left on the surface following harvest, has become gradually more popular. No-till can often increase soil carbon, soil quality and function, and reduce CO_2_ emissions when compared to conventional tilling practices (Karlen et al., 1994; Kladivko, 2001; Bolliger et al., 2006).

In the last few decades, though it varies by crop, the adoption of no-till continues to rise (Claassen et al., 2018). The most recent USDA survey estimates that no-till and strip till accounts for 45 % of total U.S. acreage in wheat (2017), 40 % in soybeans (2012), 18 % in cotton (2015), and 27 % in corn (2016) (Claassen et al., 2018). A large global meta-analysis of no-till productivity across a range of climates, soils, and crop types found that no-till management consistently out-performed conventional tillage for rainfed crops in dry climates (Pittilkow et al., 2015). However, in more mesic climates, or under irrigated conditions, differences were more variable. Some of the variation uncovered by this meta-analysis is likely due to the short duration of many of the studies – only 60 of the 520 studies lasted 10 years or longer.

Short-term research experiments are important for identifying ecosystem related changes to land management in a timely and efficient manner. In fact, for a variety of reasons, including research funding cycles, human or business constraints, and/or a need for actionable solutions, agricultural research is often performed on truncated time scales. Yet, research conducted at these shorter time scales has the potential to be misleading. Short-term research can be insufficient especially for evaluating ecosystem properties and phenomena that change slowly, such as soil structure and soil carbon, or require proper evaluation over an appropriately variable climate and pest history (Paustian et al., 1997; Rasmussen et al., 1998; Robertson et al., 2008).

Beyond the variable findings that surround the relative environmental benefits of no-till, economic concerns have also slowed the adoption of more sustainable practices (Wade and Claassen, 2017). Surveys suggest that when asked, growers responded that compared to other factors (e.g. lack of education and/or information, resistance to change, social considerations, infrastructure, or landlessness), economic concerns are the largest barrier to adopting sustainable agricultural practices, such as no-till management (Rodriguez et al., 2009). Converting a field to no-till involves both the upfront expenses of investment in novel machinery (Krause and Black, 1995), as well as the increased herbicide cost of controlling weeds, which can exceed the short-term savings associated with reduced tillage. Thus, growers may choose to avoid no-till as a result of both the uncertainty surrounding benefits as well as short-term economic hurdles.

That many of the attributes and perhaps functional benefits of no-till management may take decades to develop consistent impacts on yields, profits, and environmental outcomes begs three questions: 1) How long does continuous no-till need to be implemented (or studied) until economic and ecological benefits are consistently detectable? 2) How consistent are changes in economic and environmental attributes over long periods? 3) How many years of continuous no-till management are needed to recoup the upfront expenses of converting conventionally tilled fields to continuous no-till management? We use a 29-year experimental dataset and power analysis for a long-term research site in the upper Midwest, U.S. to 1) determine the number of years required for differences in crop yield, soil moisture, and N_2_O fluxes to be detectable, 2) investigate the consistency of trends over time, and 3) determine the number of years before continuous no-till consistently recovers initial management costs. To address questions 2 and 3, we use a moving window approach. To further address question 3, we also use a partial budgeting analysis of relative profitability.

## 2 METHODS

### 2.1 Study site and treatments

We explicitly tested the economic and ecological effects of no-till in the Main Cropping System Experiment (MCSE) of the Kellogg Biological Station Long-Term Ecological Research site (LTER) located in southwest Michigan (42°24′ N, 85°24′ W) in the northeastern portion of the U.S. Corn Belt. The mean annual air temperature at KBS is 10.1 °C, ranging from a monthly mean of −9.4 °C in January to 28.9 °C in July. Rainfall averages 1027 mm yr^−1^, evenly distributed seasonally; potential evapotranspiration exceeds precipitation for about four months of the year. Loam soils are well-drained Typic Hapludalfs developed on glacial outwash with soil carbon contents around 1% C (Syswerda et al., 2011). More details are available in Robertson and Hamilton (2015).

The MCSE was established in 1989 and includes corn (*Zea mays* L.), soybean (*Gylcine max* L.), and wheat (*Triticum aestivum* L.) rotations under varied management regimes, each replicated as 1 ha plots in six blocks. Management treatments include conventional tillage, with fertilizer and pesticides applied at rates based on soil-test recommendations, integrated pest management, and moldboard or chisel plow tillage; and no-till, similar to conventional tillage but continuous no-till (Figure 1). Nine additional non-tillage related treatments occur at the site and are not used in this study. Conventional tillage consisted of moldboard plowing in the spring from 1989 to 1998, and chisel plowing in the spring from 1999 to present. Additional tillage consisted of disking before winter wheat planting. Tillage plots were planted with a John Deere Planter. No-till management requires specialized seeding equipment such as seed drills to plant seeds into undisturbed crop residues and soil. No-till was planted with a John Deere planter drill. Both treatments were harvested using a John Deere combine. Site history prior to 1989 consisted of mixed agricultural and horticultural cropping for 100+ years, with the most recent years prior to experiment establishment dominated by conventionally managed continuous corn production. In 2009, soybean varieties were changed from conventional to transgenic (glyphosate resistant) and in 2011 corn varieties changed from conventional to transgenic (glyphosate, European corn borer, and root worm resistant), consequently reducing the expense of post planting agrichemical management. Detailed descriptions of the treatments, management protocols, and site history are provided in Robertson and Hamilton (2015).

**Figure 1:**
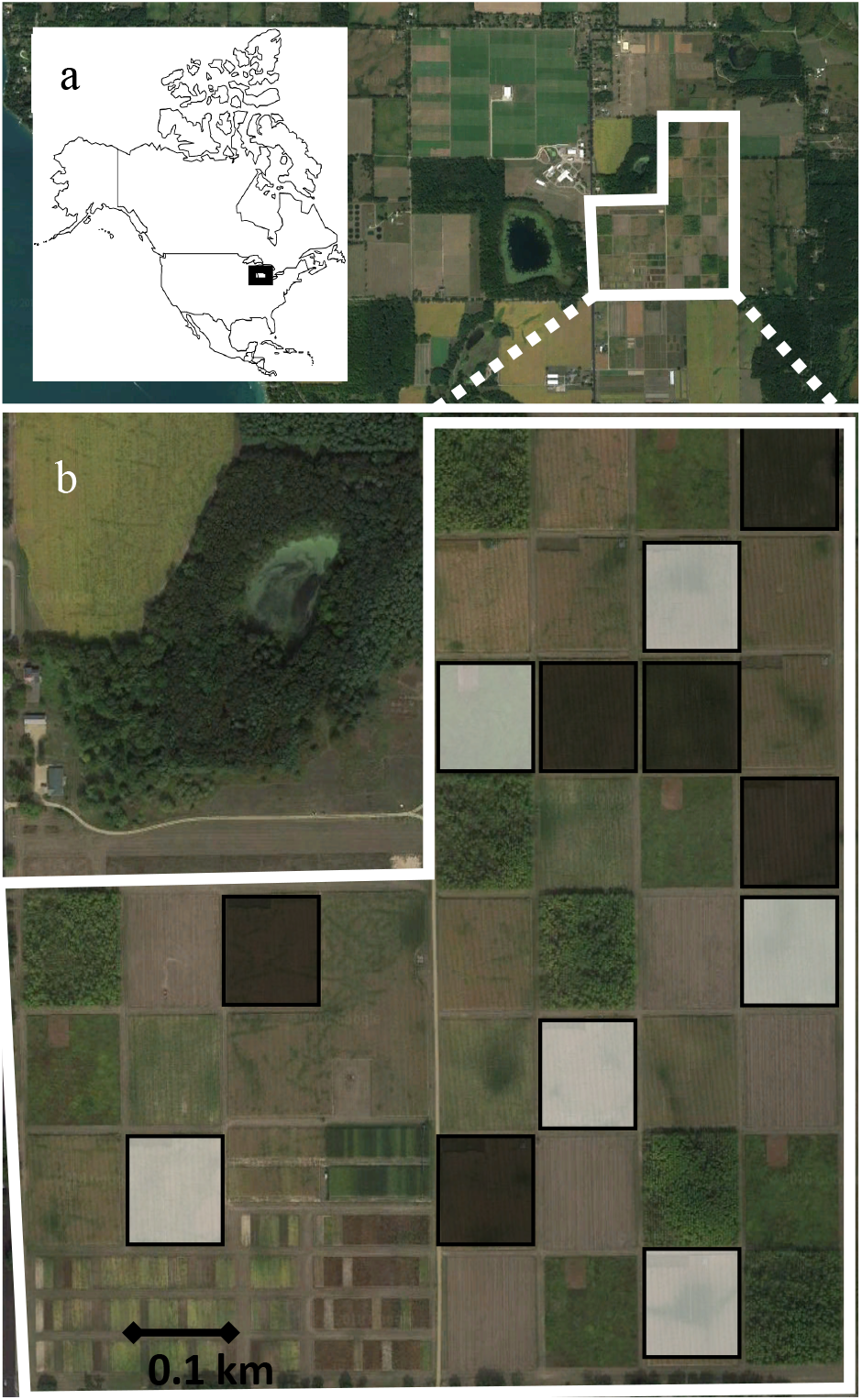
Location of study site. a) Inset of North America with outline of study region. Satellite image of the Kellogg Biological Station LTER in Hickory Corners, Michigan, U.S. b) enlarged image of Main Cropping System Experiment (MSCE). White squares show conventionally managed 1-ha blocks (N=6) and black squares show no-till managed 1-ha blocks (N=6). Blocks were established in 1988.

### 2.2 Linear regression and power analysis

To test the time required to detect the effects of tillage change (Question 1), we examined both relative profitability and environmental responses to conventional and no-till management treatments over a 29-year period from 1989 to 2017. Wheat was typically harvested in June, and soybean and corn in October and November, respectively, although some deviations occurred due to weather and crop maturity. Biomass was dried at 60° C for at least 48 h, weighed, and reported as Mg at standard moisture.

We also assessed the effect of tillage on soil moisture and N_2_O gas fluxes, our environmental response variables. Soil moisture affects microbial activity, carbon and nutrient availability and movement, and plant growth, in particular (Lal, 2004). Soil moisture was determined gravimetrically by drying 40 grams of fresh soil at 60° C for 48 hours. After drying, samples were reweighed, and gravimetric soil moisture was calculated as the difference between fresh and dry weight, expressed as dry weight (g H_2_O/g dry soil) (http://www.lter.kbs.msu.edu). Nitrous oxide (N_2_O) is an ozone depleting greenhouse gas (Ravishankara et al., 2009) and agricultural soil management is the largest anthropogenic source of N_2_O emissions globally (Paustian et al., 2016). N_2_O gas measurements were made using static chambers (Livingston and Hutchinson, 1995) at weekly to monthly intervals when soils were not frozen (Gelfand et al., 2016). Single chambers were located in four of the six blocks of each tillage treatment. Chamber lids were placed on semi-permanent aluminum bases removed only for cropping activities and accumulated headspace was sampled four times over 120 minutes. All chambers were sampled on the same dates, although no data are available for 1995. Samples were stored in 3-mL crimp-top vials and analyzed in the laboratory for N_2_O with the flux for each chamber calculated as the linear portion of the gas accumulation curve for that chamber. Nitrous oxide was analyzed by gas chromatography using a 63Ni electron capture detector. More details appear in Gelfand et al. (2016).

To test for tillage related changes in crop yield, soil moisture, and N_2_O fluxes over our 29-year period we fit linear mixed effects models to the data using crop (corn, soybean, wheat) and block (1-6 or 1-4) as fixed variables, year as a random variable, and difference in crop yield, soil moisture, and N_2_O fluxes as response variables with the ‘lmer4’ package in R (Bates et al., 2015). We then executed a power analysis for our linear mixed effects models through simulation with a traditional alpha value of 0.05, 1000 simulations, and power of 0.8, using the ‘simr’ package in R (Green et al., 2019). We also executed a second power analysis at a more liberal alpha value for comparison (alpha = 0.2, a lower, less confident level of significance).

### 2.3 Moving window

To investigate our second question, concerning the variability of trends in our long-term dataset, we used the same response variables described above and a moving window approach. Conceptually, this provides a trajectory of the relationship of each response variable with time, and describes how the fit of linear mixed effects models results vary with different sample periods, start years, and durations. We used a moving window algorithm developed in R (Bahlai, 2019) to measure the overall trajectory and consistency of our response variables throughout our 29-year dataset. First, we fit linear mixed effects models to defined subsets of data and produced summary statistics of interest (e.g. slope of the relationship between the response variable and time, standard error of this relationship, and p-value). The moving window then iterated through the entire dataset at set intervals. We used moving windows of three-year periods or longer, fed each interval through the algorithm described above, and compiled resulting summary statistics. The direction and magnitude of statistically significant slopes are plotted against corresponding window length (number of years) to investigate the relationship between trend consistency, direction, and magnitude with study duration.

### 2.4 Partial Budgeting Relative Profitability Analysis

To answer our third question, comparing the expense of implementation and management of the two tillage systems, we used a partial budgeting relative profitability analysis combined with the moving window approach, described above. We determined the relative profitability benefits of no-till management as the difference in annual gross margins between no-till and conventional till treatments (Cimmyt and Cimmyt, 1988). Gross margins were calculated as annual crop grain yield multiplied by current year crop price ($/kg) minus costs that varied between the two treatments. We subdivided the gross margins by tracking differences in crop revenues and expenses between the treatments. For revenues, we determined differences in yield as described in section 2.1 and crop prices as the U.S. monthly average price received for corn, soybean, and wheat at time of harvest (December, November, and June, respectively) (FarmDoc, 2019) (ESM Table 1).

**Table 1:**
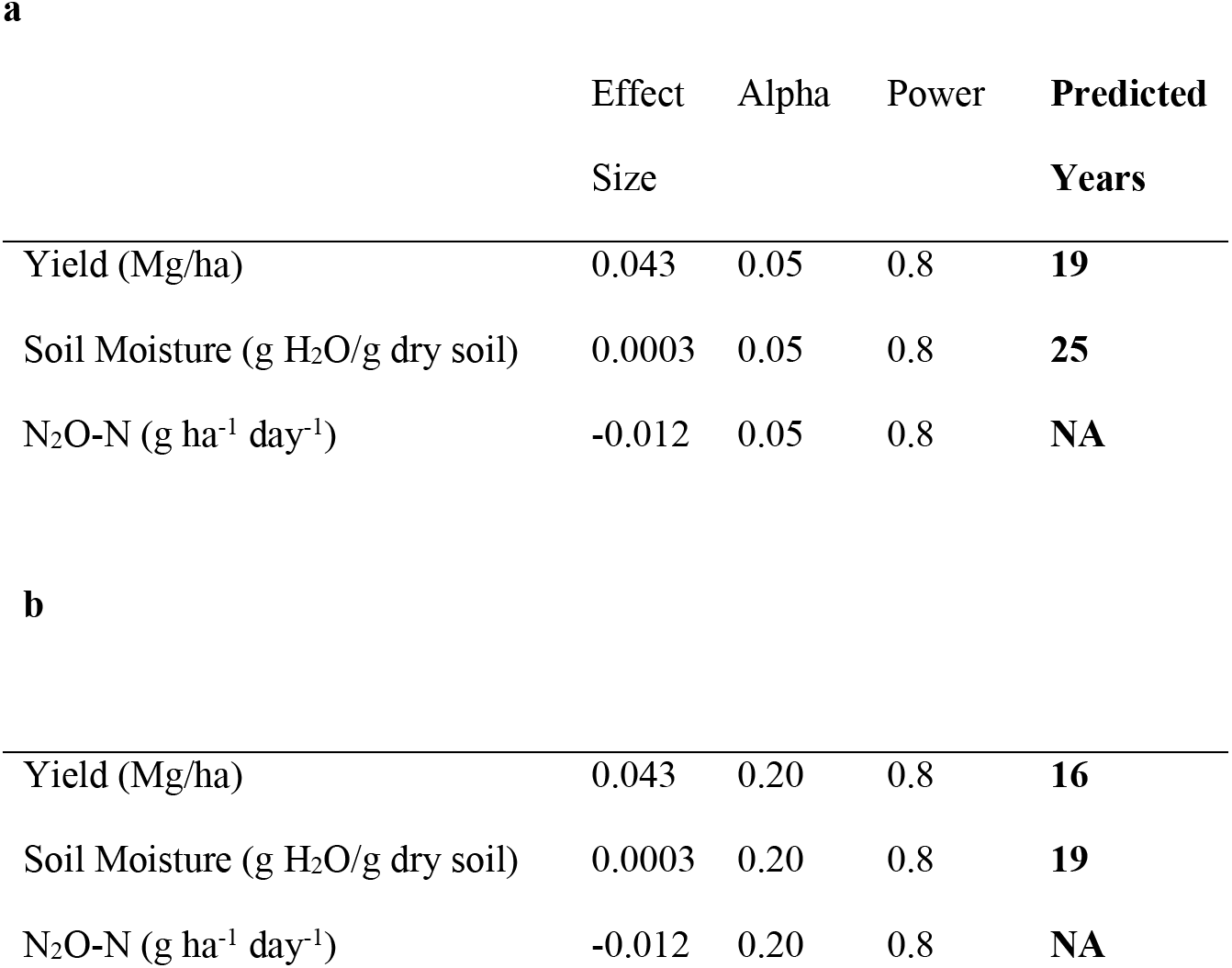
Summary of a) power analysis predicting the number of years needed to detect effects of no-till treatment on crop yield (Mg/ha), soil moisture (g H_2_O/g dry soil), and N_2_O-N (g ha^−1^ day ^−1^) in linear mixed effects models given a conservative alpha value of 0.05 and power of 0.8, and b) for a more liberal alpha value of 0.2 and power of 0.8.

|  | Effect<br>Size | Alpha | Power | **Predicted**<br>**Years** |
| --- | --- | --- | --- | --- |
| Yield (Mg/ha) | 0.043 | 0.05 | 0.8 | **19** |
| Soil Moisture (g H<sub>2</sub>O/g dry soil) | 0.0003 | 0.05 | 0.8 | **25** |
| N<sub>2</sub>O-N (g ha<sup>-1</sup> day<sup>-1</sup>) | -0.012 | 0.05 | 0.8 | **NA** |
| **b** |  |  |  |  |
| Yield (Mg/ha) | 0.043 | 0.20 | 0.8 | **16** |
| Soil Moisture (g H<sub>2</sub>O/g dry soil) | 0.0003 | 0.20 | 0.8 | **19** |
| N<sub>2</sub>O-N (g ha<sup>-1</sup> day<sup>-1</sup>) | -0.012 | 0.20 | 0.8 | **NA** |

We determined the relative expenses of no-till management as the combined difference in expense of input and custom work rates between the two treatments. We determined input expenses only for those chemical inputs that differed in application between the two treatments as described in the KBS ‘Expanded Agronomic Log’ (http://www.lter.kbs.msu.edu). All differences were first converted to fluid oz/ha or kg/ha and then, using historic prices (USDA, 2019), into differences in expense and revenue, expressed in $/ha each year. We determined differences in custom work rates between treatments using information from the Michigan State University Department of Agricultural Food and Resource Economics (Michigan State University, 2019) between 1994 and 2018. Years prior to 1994 and years with missing values were extrapolated and interpolated, respectively, from an estimated linear relationship. Conventional tillage expenses included the custom rates for tillage (moldboard or chisel plow), soil finishing, and planting. No-till expenses included custom rates for planting, spraying, and mowing. Custom work estimates include the expense of labor, fuel, and equipment rental. Custom rates in Michigan were compared to those calculated in Iowa between 1995-2014 (Edwards and Johanns, 2014), and were found to follow similar trajectories over time. Finally, differences in revenue ($/ha) and expense ($/ha) were converted from different time periods to present values (base year 2017) using the present value formula: *PV*=*FV*(1+*i*) ^*n*^, where “*PV*” is present value, “*FV*” is future value, “*i*” is interest rate (we used 5%), and “*n*” is the number of years. Derivation of value estimates of expenses and revenue by year are given in supplemental materials (ESM Table 1). We determined accumulated expenses, revenue, and difference between treatments (no-till – conventional) to determine when no-till managed blocks recuperate initial implementation and ongoing maintenance expenses. Curves were estimated using third order polynomials: USD/ha = a*Year + b* Year ^2^ + c* Year ^3^, where ‘a’, ‘b’, and ‘c’ are coefficients (estimates provided in ESM Table 2).

**Table 2:** Trend summaries for crop yield (Mg/ha), gravimetric soil moisture (g H_2_O/g dry soil), and N_2_O-N (g ha^−1^ d-^1^) for each block from moving window analysis applied to linear mixed effects models.

|  | Total<br>Trends | Total<br>Significant<br>Trends | Positive<br>Trends | Positive<br>Significant<br>Trends | Negative<br>Trends | Negative<br>Significant<br>Trends |
| --- | --- | --- | --- | --- | --- | --- |
| Yield (Mg/ha) | 379 | 182 | 250 | 170 | 129 | 12 |
| Soil Moisture (g H<sub>2</sub>O/g dry soil) | 379 | 139 | 299 | 128 | 80 | 11 |
| N<sub>2</sub>O-N (g ha<sup>-1</sup> day<sup>-1</sup>) | 142 | 10 | 65 | 3 | 77 | 7 |

To understand the consistency of relative profitability over time intervals, we applied the moving window approach as described above. Conceptually, we are interested in the trajectory of the relationship between accumulated relative profitability between no-till and conventional agriculture with time, and how linear mixed effects model outputs vary with the starting point of sample period and sample period duration. The direction and magnitude of statistically significant slopes were plotted against corresponding window length (number of years) to investigate the relationship between relative profitability trend consistency, direction, and magnitude with study duration.

## 3 RESULTS

### 3.1 Summary of study site and treatments

Between 1989 and 2017, differences in corn yield between treatments (no-till – conventional) ranged from −3.57 Mg/ha in 1999 to 2.91 Mg/ha in 2008 and averaged 0.68 Mg/ha (SE: 0.017). Differences in soybean yield (no-till – conventional) ranged from −2.68 Mg/ha in 1994 to 1.07 Mg/ha in 2006 and averaged 0.306 Mg/ha (SE: 0.007). Differences in wheat yield (no-till – conventional) ranged from −4.14 Mg/ha in 2001 to 1.62 Mg/ha in 2010 and averaged 0.042 Mg/ha (SE: 0.001). Over the same time period, differences in soil moisture between the two treatments across crop types (no-till – conventional) ranged from −0.026 g H_2_O/g dry soil in 1990 to 0.05 g H_2_O/g dry soil in 2013 and averaged 0.02 g H_2_O/g dry soil (SE: 0.002). Differences in N_2_O gas fluxes between the two treatments (no-till – conventional) ranged from −4.3 g N_2_O-N ha^−1^ d^−1^ in 2007 to 6.9 g ha^−1^ day^−1^ in 2000 and averaged 0.4 g ha^−1^ d^−1^ (SE: 0.2).

### 3.2 Linear regression and power analysis

Using linear mixed effects models to address our first question, we found a significant positive relationship between window duration and difference in crop yield (Estimate: 0.043, T: 4.96, P < 0.001) (Figure 2a). Further, we found a significant positive relationship between differences in soil moisture and duration between no-till and conventionally managed treatments (Estimate 0.0003, T: 4.005, P < 0.001) (Figure 2b). Lastly, we found a non-significant relationship between differences in N_2_O fluxes and window duration between no-till and conventionally managed treatments (Estimate: −0.012, T: −0.39, P: 0.70) (Figure 2c).

**Figure 2:**
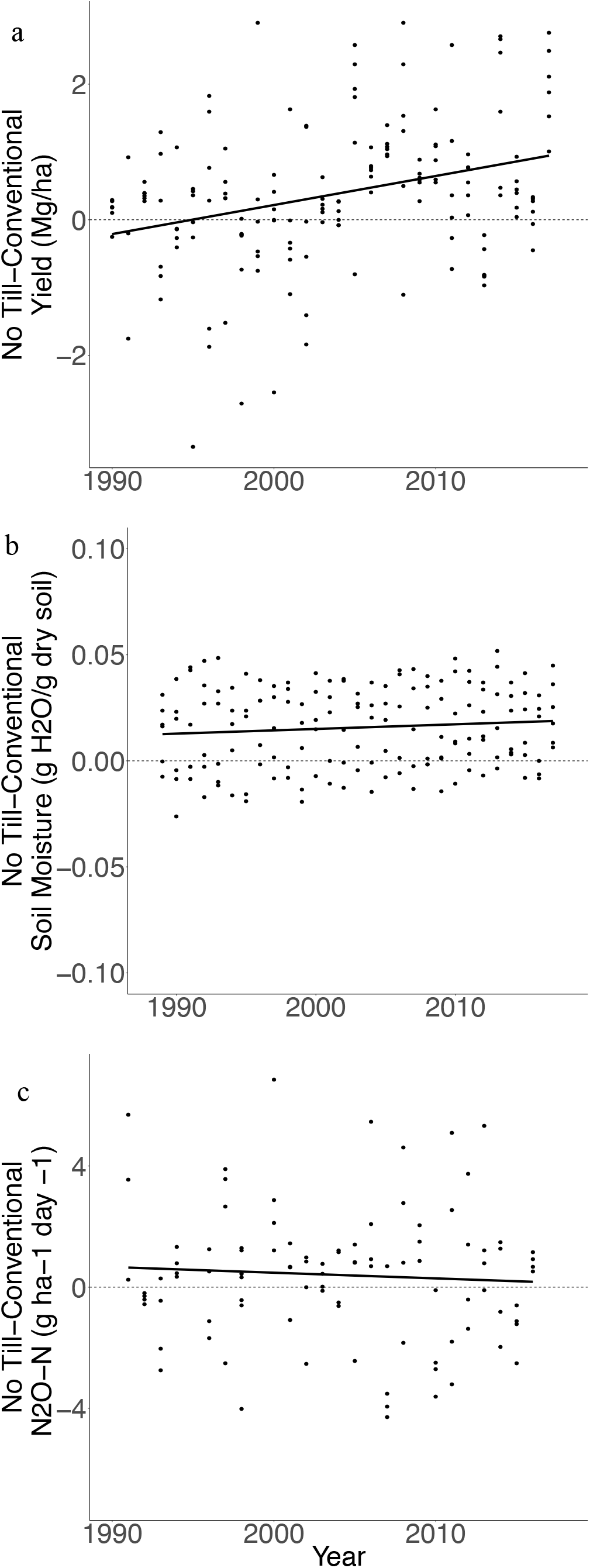
Scatterplots showing a) the difference in crop yield (Mg/ha) between no-till and conventional treatments within each block and year, b) the difference in gravimetric soil moisture (g H_2_O/g dry soil) between no-till and conventional treatments within each block and year, and c) the difference in N_2_O-N (g ha^−1^ day^−1^) between no-till and conventional treatments within

In a conservative power analysis, we found that with a calculated effect size of 0.043, alpha of 0.05, and power of 0.8, it would take at least 19 years to detect a significant effect of the no-till treatment on crop yield using a linear mixed effects model. Likewise, with a calculated effect size of 0.0003, it would take 25 years to detect a significant effect of the no-till treatment on soil moisture. Lastly, even with a calculated effect size of −0.012, we would not see a significant effect of the no-till treatment on N_2_O flux (Table 1a). Using a more liberal alpha value that better represents the interests of growers (alpha = 0.2) we see that, given our effect sizes, it would take 16 and 19 years to detect significant effects of no-till management on crop yield and soil moisture, respectively (Table 1b).

### 3.3 Moving window

The moving window algorithm formed 379, 379, and 142 windows of lengths between three and 29 years for our long-term datasets (difference in yield, soil moisture, and N_2_O fluxes, respectively). Addressing our second question concerning the consistency of trends, we found that for differences in yield, 250 windows had a positive slope (66%) and 129 windows had a negative slope (34%). Of the positive trending windows, 170 were significant at the alpha <0.05 level; of the negative trending windows, 12 were significant. (Figure 3a, Table 2). For soil moisture, we found that 299 windows had a positive slope (79%) and 80 windows had a negative slope (21%). Of the positive trending windows, 128 were significant; of the negative trending windows, 11 were significant (Figure 3b, Table 2). Lastly, for N_2_O fluxes, of the 142 windows, we found 65 windows were positively trending (46%), 77 were negatively trending (54%), and three of the positive windows and seven of the negative windows were found to be significant (Figure 3c, Table 2).

**Figure 3.**
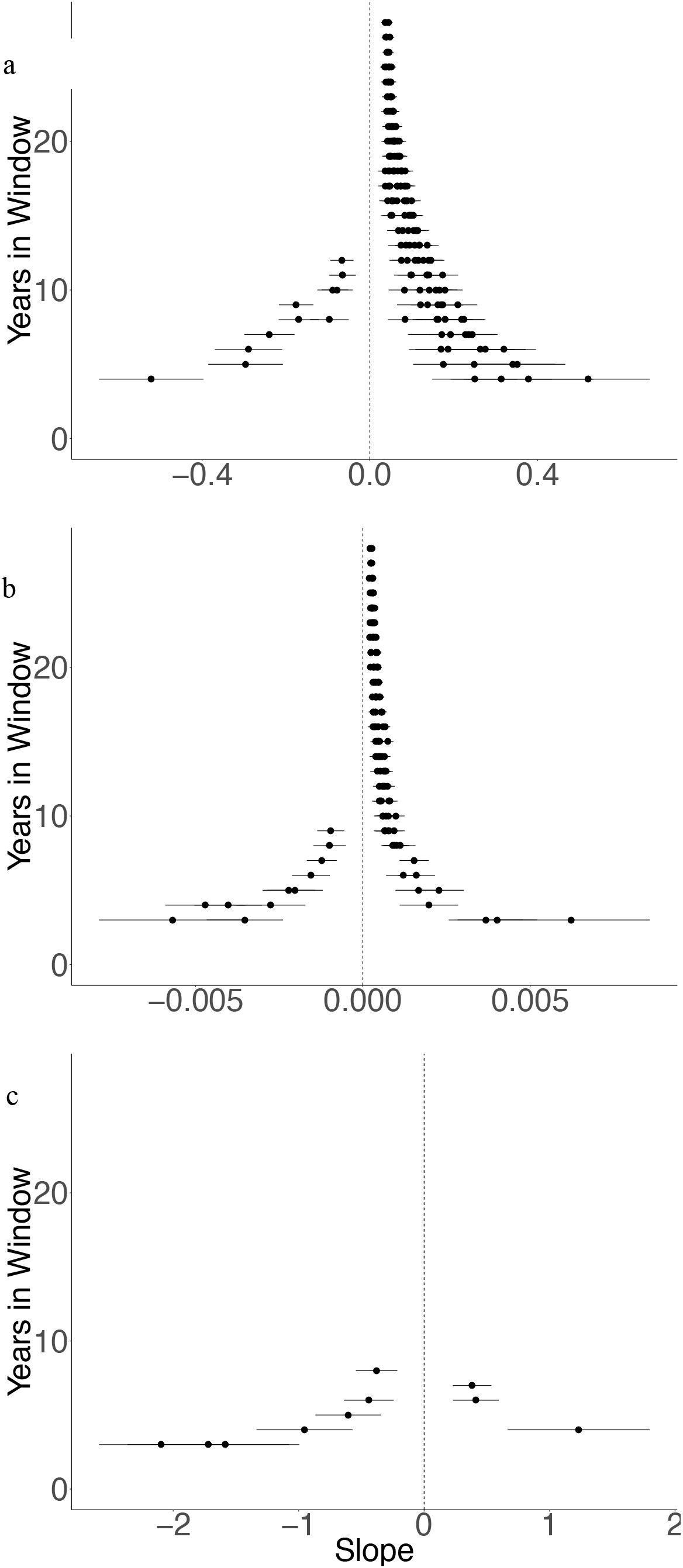
Results from the moving window analysis. Scatterplots showing the relationship between slope and the number of years in a given window for a) crop yield (Mg/ha), b) gravimetric soil moisture (g H_2_O/g dry soil), and c) N_2_O-N (g ha^−1^ d^−1^). Slopes were determined using linear mixed effects models using crop (corn, soybean, wheat) and block (1-6) as fixed variables, and year as a random variable for each window of time three years in length or longer, between 1988 and 2017. Only models with statistically significant slopes at the alpha < 0.05 level are shown. Dots represent slope and solid lines represent standard error. Positive slopes indicate an accruing benefit of no-till management over time when compared directly to conventionally managed blocks. The vertical black dotted line indicates a slope of zero, where the benefits of no-till management have saturated.

### 3.4 Partial Budgeting Relative Profitability Analysis

Because the relative cost of no-till flattened out over time while crop yields under no-till increasingly gained over conventional tillage, the accumulated expenses ($/ha) of no-till compared to conventional management did not grow as fast as the accumulated revenue. Hence, gross margin differences reveal that while no-till was a money loser in early years, by the end of 2002 (13 years after implementation), no-till recuperated initial losses at our site. By 2017, no-till had accumulated nearly $2,000/ha in differences in relative profitability as measured by partial budgets (Figure 4, ESM Table 1, ESM Table 2). Using the moving window algorithm on the partial budgeting analysis of relative profitability, we see that of the 379 windows formed, 338 were significant at the p < 0.05 level. While most trends were positive, of windows between 3 and 10 years long, 37% (57/156) were negatively trending, 19% (26/137) windows between 11 and 20 years long were negatively trending, and of the windows between 21-29 years long none (0/46) were negatively trending (Figure 5, Table 3).

**Table 3:** Trend summaries for partial budgeting analysis of relative profitability comparing the expense of implementation and management of the two tillage systems from moving window analysis applied to a linear mixed effects model using crop (corn, soybean, wheat) and block (1-6) as fixed variables, and year as a random variable.

|  | Total<br>Trends | Total<br>Significant<br>Trends | Positive<br>Trends | Positive<br>Significant<br>Trends | Negative<br>Trends | Negative<br>Significant<br>Trends |
| --- | --- | --- | --- | --- | --- | --- |
| Early (3-10 yrs) | 188 | 156 | 115 | 99 | 73 | 57 |
| Mid (11-20 yrs) | 145 | 136 | 115 | 26 | 30 | 110 |
| Long (21-29 yrs) | 46 | 46 | 46 | 46 | 0 | 0 |
| Full (3-29 yrs) | 379 | 338 | 276 | 255 | 103 | 83 |

**Figure 4.**
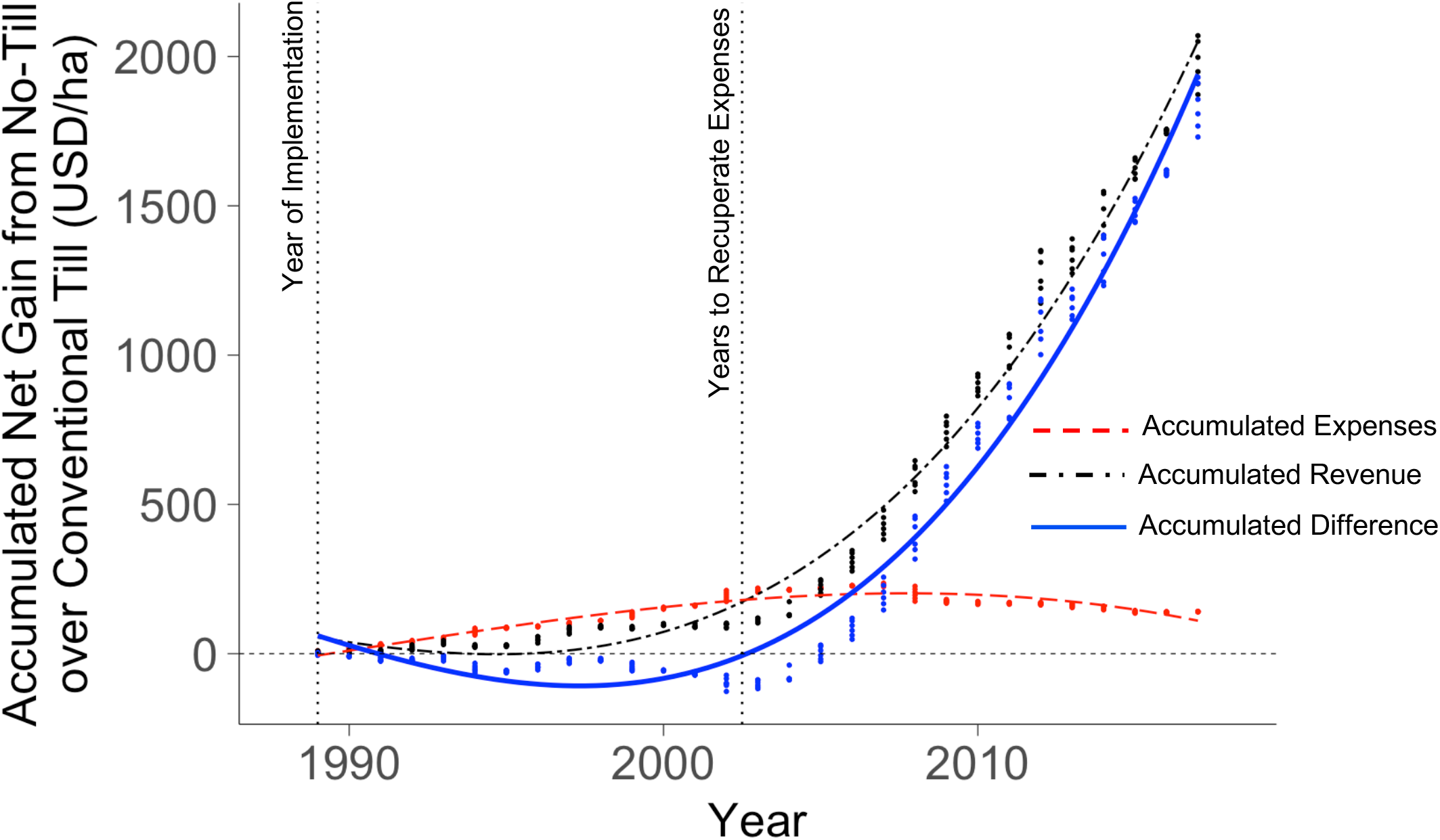
Partial budgeting analysis of relative profitability plot comparing the expense of implementation and management of the two tillage systems. Dots represent values for each block each year. Accumulated expenses, shown as red dots and the dashed red line, include differences in custom hire ($/ha) and input expenses ($/ha) between the two treatments (no-till – conventional). Accumulated revenue, shown as black dots and the dashed black line, include the difference in yield between the treatments (no-till – conventional) (Mg/ha) each year and historic crop prices at time of harvest ($/kg). The blue dots and blue line describe accumulated net gain from no-till over conventionally tilled management. Values were converted from different time periods to present values using the present value formula: *PV*=*FV*(1+*i*) ^*n*^, where “*PV*” is present value, “*FV*” is future value, “*i*” is interest rate (we used 5%), and “*n*” is the number of years. The vertical, black dotted line estimates the time at which accumulated expenses equal accumulated revenue

**Figure 5.**
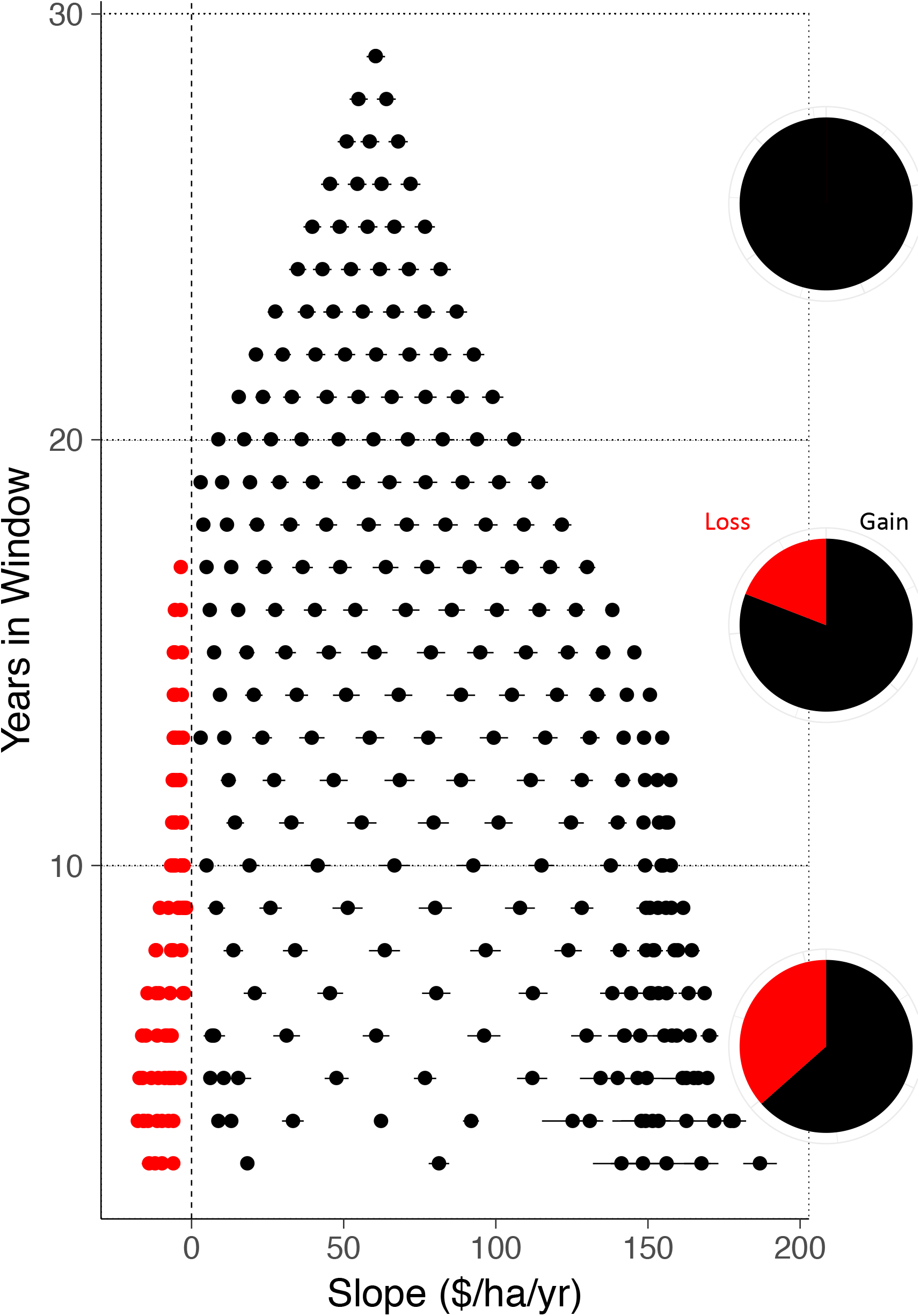
Scatterplot showing the relationship between slope ($/ha/yr) and the number of years in a given window in terms of accumulated difference between no-till and conventional management ($/ha). Slopes were determined using linear mixed effects models using crop (corn, soybean, wheat) and block (1-6) as fixed variables, year as a random variable, and accumulated gain in no-till over conventional till for each window of time three years in length or longer, between 1989 and 2017. Only models with statistically significant slopes at the alpha < 0.05 level are shown. Dots represent slope and solid lines represent standard error. Positive slopes, shown in black, indicate accruing gain in relative profitability of continuous no-till management when compared directly to conventionally managed blocks. Negative slopes, shown in red, indicate accruing loss of no-till management over time when compared directly to conventionally managed blocks. Pie charts show the percent of trends that are significantly negative (shown in red) and significantly positive (black) for early windows (0-10 years long, bottom), middle windows (11-20 years, middle), and long windows (21-29 years, top).

## 4 DISCUSSION

Using a long-term experimental dataset spanning nearly 30 years, combined with power analysis and a moving window approach, we found that both yield and soil moisture require periods 15 years or longer to generate consistent results in this rainfed corn-soybean-wheat system in the upper Midwest U.S. In fact, given our effect sizes, it would take 16 and 19 years to detect significant effects of no-till management on crop yield and soil moisture, respectively, even with a liberal alpha value (alpha = 0.2). Analyses performed on periods shorter than 15 years suggest misleading trends, even though these findings were sometimes statistically significant.

Through relative profitability analysis, we found that 13 years were needed to fully recover the initial expenses of no-till management in our system, and that the longer the implementation of no-till, the greater likelihood of relative profitability, regardless of stochastic effects. Here, we show that more than a decade is needed to detect the consistent benefits of continuous no-till on these economic and environmentally important attributes, suggesting that a shorter-term experiment could have led to contradictory, and potentially misleading results.

Nine out of 45 significant results for periods shorter than 10 years support a negative relationship between no-till and the difference in yield over time (20%), in direct contradiction to the positive pattern we observed. Only analyses performed on periods 10 years or longer resemble more closely the broader positive pattern. Further, 31% (11/35) of the analyses on periods shorter than 10 years indicate a statistically significant negative relationship for soil moisture over time, in direct contradiction to the relationship suggested by longer periods. Among the periods tested for N_2_O fluxes, 10 of 142 were found to contain a statistically significant trend (seven negative and three positive). This low detection of significant trends is consistent with recent N_2_O flux meta-analyses (e.g., Van Kessel et al., 2013), where only for studies greater than 10 years duration, particularly in drier climates, were trends of lower fluxes in no-till than conventional management significant. Results from the present site suggest that such a trend may take substantially longer (perhaps scores of decades) to detect, likely in part due to relative increases in N_2_O fluxes for the rotation’s corn and soybean years offsetting decreases in wheat years (Gelfand et al., 2016). Trends may not emerge until the rotation or some other important aspect of agricultural management changes, such as fertilizer rate (Shcherbak et al., 2014). Nevertheless, no-till does not (yet) have consistent detectable effects on N_2_O fluxes at this site, as noted for earlier shorter-term analyses (Robertson et al., 2000; Grandy et al., 2006; Gelfand et al., 2016). The importance of N_2_O-N for mitigating the global warming impact of intensive cropping systems underscores the importance of better understanding such time-dependent trends (Six et al., 2004).

Lastly, we show that longer periods of implementation increase the likelihood that continuous no-till agriculture becomes more profitable than tillage. Immediately following implementation of no-till management (periods between 3 and 10 years) more than one third of periods resulted in the loss of net revenue (37%). However, as periods became longer, the likelihood of greater relative profitability increased. In fact, for periods between 11 and 20 years long, fewer than 20% of periods lost revenue, and in periods between 21 and 29 years, all periods were profitable. Overall, 86% of periods greater than 10 years were profitable. We also note that, beyond the purely fiscal benefits accrued, were value given for environmental impacts of continuous no-till (i.e. carbon sequestration, reduced nitrate leaching, etc.) the benefits would further increase.

Our results show that even in the absence of an overarching trend, spurious relationships in temporal processes or short-term trends associated with stochastic processes are common in tillage systems. Thus, the conflicting trends and predictions noted in previous studies concerning the impact of tillage on crop yield, soil moisture, and N_2_O fluxes may be explained, at least in part, by their durations: strong, statistically significant relationships between parameters and duration may have been the result of high variation in the system over shorter time periods. This phenomenon of confident though misleading results highlights the importance of long-term studies for detecting trends and informing management recommendations with confidence.

To maximize the impact of research at any time scale, it is essential to understand how patterns emerge as studies become longer, enabling us to more effectively extrapolate results to long-term patterns. As our results show, variation can be highly idiosyncratic and dependent on study duration. Predicting the future effects of continuous no-till depends on understanding both the short-and long-term dynamics of crops following significant changes in management. This is likely to be especially important for detecting the consequences of slow to change properties like soil organic matter accretion following no-till initiation.

Our results highlight the importance of not only study duration, but also of the selection of study starting and ending points. If a study period captures an outlying data point in a system’s natural variability near the beginning or end of the study, those years are likely to be disproportionately influential on statistical outcomes and thus on conclusions, and presumed management implications (Swinton and King, 1991; White, 2019). Periods that reveal contrary results may be the response of high variation, possibly caused by extreme weather events, changes in crops, or other system level idiosyncrasies.

In the case of no-till implementation, transient dynamics leading to short-term risks associated with no-till management are likely to be captured by short-term studies that focus on the establishment of no-till at new locales. These risks include increased pest and disease danger, altered nitrogen cycling, and increased nutrient requirements due to nutrient immobilization under cooler soil temperatures (Baker et al., 1996). Also, due to the potential slower soil warm-up in the spring, no-till management may result in stunted growth in initial years. Lastly, increased pressure by herbicide resistant weeds may cause future problems (Van Deynze et al., 2018).

However, as revealed by our results, benefits at our site accrue over time. As Choudhary and Baker (1994) predicted, despite potential negative results in the first few years of no-till, benefits of the reduced fertilizer requirements and pest protection, as well as an increased stable crop yield, are only realized with long-term management. Further, because continuous no-till can be economically attractive for other reasons in the long term (e.g. reduced machinery fuel, energy, and maintenance costs (Lal et al., 2007a; Rathke et al., 2007)), our results are consistent with recommendations to support the long-term adoption of no-till management despite initial losses.

## 5 CONCLUSIONS

We used 29 years of no-till crop management data to reveal the temporal processes and long-term impacts of a change in agricultural management at a site in the U.S. corn belt. We illustrate that management recommendations based on short term studies can be contradictory because spurious, misleading trends can appear in time series at rates between 20 and 50% of the time, even independent of stochastic elements associated with external disturbances. Furthermore, the initiation of a new experiment almost certainly represents a strong disturbance to an ecosystem, thus the early years in a study involving temporal processes may produce data that is not representative of the system’s equilibrium behavior. Our results are consistent with recommendations to support the long-term adoption of no-till agricultural management despite initial losses.

## Supporting information

ESM 1

ESM Table 2

## 6 ACKNOWLEDGMENTS

This work was performed on the traditional Anishinaabe land where Hickory Corners, Michigan is currently located. Support for this work was provided by the National Science Foundation Long-term Ecological Research Program (DEB 1832042) at the Kellogg Biological Station, USDA National Institute on Food and Agriculture, and by Michigan State University AgBioResearch. Thanks to KBS colleagues for providing data and maintaining experimental infrastructure, especially Sven Bohm for database curation. The scaling algorithm was developed by CB in response to conversations with Elise Zipkin, Ilya Gelfand, Kaitlin Stack Whitney, Easton White, and Julia Perrone, and is funded by the National Science Foundation Directorate for Computer and Information Science and Engineering (OAC 1838807).

