## Supplementary material for "Long-term research needed to avoid spurious and misleading trends in sustainability attributes of no-till": ESM Table 2

ESM Table 2: Model summaries for third order polynomials for accumulated expenses, revenue, and difference between treatments (no-till – conventional). Values were converted from different time periods to present values using the present value formula: 𝑃𝑉=𝐹(1+𝑖) ^𝑛^, where “*PV*” is present value, “*FV*” is future value, “*i*” is interest rate (we used 5%), and “*n*” is the number of years.

| Accumulated Difference in Expense ($/ha) | Estimate | Std. Error | t value | p value |
| --- | --- | --- | --- | --- |
| Intercept | 136.79 | 1.39 | 98.14 | <0.001 |
| Year | 589.51 | 18.39 | 32.06 | <0.001 |
| Year^2^ | -552.72 | 18.39 | -30.06 | <0.001 |
| Year^3^ | -101.66 | 18.39 | -5.53 | <0.001 |

| Accumulated Difference in Revenue ($/ha) | Estimate | Std. Error | t value | p value |
| --- | --- | --- | --- | --- |
| Intercept | 504.14 | 5.54 | 91.00 | <0.001 |
| Year | 7201.85 | 73.08 | 98.56 | <0.001 |
| Year^2^ | 3587.37 | 73.08 | 49.09 | <0.001 |
| Year^3^ | 518.34 | 73.08 | 7.093 | <0.001 |

| Accumulated Difference in Revenue - Expense ($/ha) | Estimate | Std. Error | t value | p value |
| --- | --- | --- | --- | --- |
| Intercept | 367.35 | 6.74 | 54.52 | <0.001 |
| Year | 6612.34 | 88.87 | 74.40 | <0.001 |
| Year^2^ | 4140.08 | 88.87 | 46.59 | <0.001 |
| Year^3^ | 620.00 | 88.87 | 6.98 | <0.001 |
